# Characterization of *spe-40/Fam187* identifies a deeply conserved sperm protein at the *C. elegans* fertilization synapse

**DOI:** 10.64898/2026.05.14.723898

**Authors:** Jeyashree Nathan Elango, Isaac H Shin, Anushree Gurjar, Amber R Krauchunas

## Abstract

Fertilization is the process in which two specialized cells, the sperm and egg, interact, adhere, and fuse their membranes. This occurs in all sexually reproducing organisms. Several transmembrane and secreted proteins have been shown to be required for fertilization. Genetic mutations can alter these proteins and disrupt fertilization, leading to reduced or no offspring. When fertilization-specific sperm proteins are mutated, sperm production, motility, and activation are unaffected, but the sperm lose the ability to successfully fertilize an egg. In this study, we report on the sperm-specific protein SPE-40/FAM187, which is a single-pass transmembrane protein with an immunoglobulin-like domain. When *spe-40* is mutated in *C. elegans* the animals are severely sub-fertile due to a sperm-specific defect. All the characteristics of the sperm that we have evaluated in the mutant are normal, yet sperm lacking SPE-40 do not fertilize. SPE-40 has orthologs in other species, including humans. Thus, we have established a role for SPE-40/FAM187 in fertilization that suggests it represents a conserved component of the fertilization synapse.

## INTRODUCTION

Fertilization is a cell fusion event mediated through proteins expressed by sperm and egg that recognize, adhere and ultimately form a single cell zygote. Fertilization-competent sperm display proteins on their surface that recognize and interact with the oocyte.^1,2^ However, the proteins required for fertilization are not fully understood. Identifying the proteins responsible for fertilization will unravel the fertilization synapse and decode its mechanistic model.

Studies in *C. elegans* have identified nine sperm-specific proteins whose sole purpose is to mediate fertilization.^3–12^ Absence of any one of these proteins results in complete or near complete sterility. Even though the sterility is due to a sperm-specific defect, mutations in the genes that encode these proteins result in no observable defects in sperm production, morphology, activation, or competition. Only two of these *C. elegans* genes have clear orthologs in other species; *spe-49* and *spe-42* have orthologs *Dcst1* and *Dcst2* in vertebrates and *sneaky* in Drosophila.^13–15^ The *C. elegans* SPE-45 protein also has the same domain structure as vertebrate IZUMO1 suggesting an evolutionarily conserved relationship.^7,8^

In this study, we characterize *spe-40* mutants and show that SPE-40 is a sperm protein specifically required for mediating fertilization in *C. elegans*. Originally isolated in a forward genetic screen, we establish that a mutation in *Y37E11AR*.*7* is responsible for *spe-40* mutant phenotype. The *spe-40* gene is homologous to *Fam187* in vertebrates. The discovery of another evolutionarily conserved fertilization protein suggests that SPE-40/FAM187 may be a core component of the fertilization machinery.

## RESULTS

### SPE-40 mutants are severely sub fertile due to a sperm specific defect

Unmated *spe-40(it133)* hermaphrodites produce mostly unfertilized oocytes, averaging only 2.6 live progeny, in contrast to wild-type hermaphrodites that produce an average of 305 live progeny (Fig 1A). However, when mated with fertile males, the fertility of *spe-40(it133)* hermaphrodites can be restored by the presence of wild-type sperm. As a control we mated fertile males to feminized (*fog-2*) hermaphrodites. *fog-2* hermaphrodites do not produce any sperm, but have no other somatic or reproductive defects.^16^ Because we observe no significant difference between the number of progeny produced by *spe-40(it133)* hermaphrodites mated to fertile males and *fog-2* hermaphrodites mated to fertile males (p=0.17), we conclude that the reduction in *spe-40(it133)* self-fertility is due to a sperm-specific defect. To test if *spe-40* also affects male fertility, we mated *spe-40(it133)* males with *fog-2* hermaphrodites. We found that the *spe-40(it133)* males also exhibit reduced progeny production, averaging only 3.7 cross progeny (Fig 1A). Thus, we conclude that the sterile phenotype of *spe-40* mutants is due to a sperm-specific defect and that *spe-40* is required in both hermaphrodite-derived and male-derived sperm.

**Figure 1:**
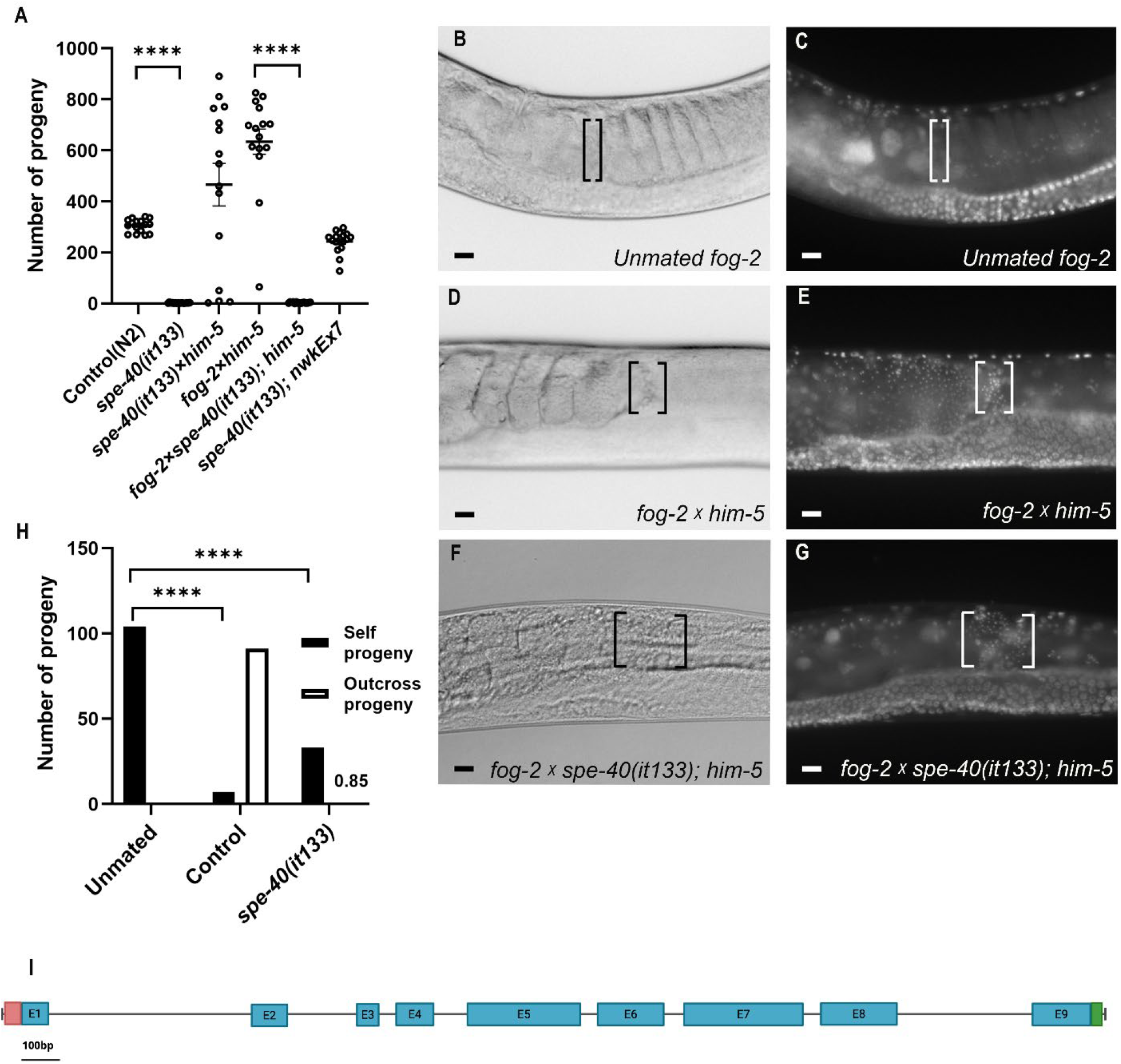
*spe-40* is required for hermaphrodite and male fertility, but has no effect on sperm transfer, sperm migration to the spermatheca, or sperm competition. **(A)** Brood sizes of *spe-40* hermaphrodites and males compared to controls. N≥15, **** indicates p<0.0001 **(B-G)** DIC imaging and DAPI staining of *fog-2* hermaphrodites. The spermatheca is indicated by brackets, the oviduct is to the right of the brackets and the uterus is to the left of the brackets. Scale bar = 12.5 µm **(H)** Sperm competition assay showing unmated *dpy-5* hermaphrodites, *dpy-5* hermaphrodites crossed with *him-5* males (Control), and *dpy-5* hermaphrodites crossed with *spe-40(it133); him-5* males. N≥15, **** indicates p<0.0001 **(E)** Schematic representation of *spe-40* gene showing 9 exons (blue bars), 5’ UTR (red) and 3’ UTR (green).

### Sperm from *spe-40* mutants are able to migrate to and maintain their position in the spermatheca and outcompete hermaphrodite self-sperm

When *C. elegans* mate, sperm from the male are transferred to the uterus of the hermaphrodite and they must migrate through the uterus to the spermathecae where fertilization occurs.^17^ To identify if sperm from *spe-40* mutant males can reach and remain in the spermathecae of hermaphrodites, *spe-40(it133)* males were crossed with *fog-*2 hermaphrodites and the hermaphrodites were then fixed and stained with DAPI. When stained with DAPI the sperm nuclei appear as small bright puncta. While unmated *fog-2* hermaphrodites have no sperm (Fig 1B-C), we observed that both control and *spe-40(it133)* males transfer sperm and the sperm migrate to the site of fertilization (i.e. the spermathecae) (Fig 1D-G).

Sperm from *C. elegans* males outcompete hermaphrodite self-sperm in fertilizing the egg.^17–20^ Since sperm from *spe-40* mutant males reach the site of fertilization, we tested whether they retain their competitive advantage. We crossed *spe-40(it133)* males with *dpy-5* hermaphrodites which have wild-type self-sperm. The recessive *dpy-5* mutation allows us to distinguish between self-progeny and outcross progeny; self-progeny are homozygous for the *dpy-5* mutation and have a visible Dpy phenotype while outcross progeny are heterozygous and appear phenotypically wild-type. Unmated *dpy-5* hermaphrodites produce only self-progeny. When mated to *him-5* males, sperm from the males fertilize the majority of the eggs and the number of self-progeny is reduced. When *dpy-5* hermaphrodites are mated with *spe-40* males, we also observe a significant reduction in the number of self-progeny compared to unmated hermaphrodites (Fig 1H). This indicates that sperm from *spe-40* mutants can still outcompete hermaphrodite self-sperm despite their failure to produce outcross progeny.

### Sperm lacking SPE-40 have normal morphology and activation but do not fertilize

For every fertilization event, the number of stored sperm in the spermathecae is reduced by one. Since *C. elegans* hermaphrodites only produce sperm during the final larval stage, adults have a finite number of self-sperm available for fertilization. Once all sperm are utilized, the worm will start producing unfertilized oocytes. DAPI staining of day one adults confirmed the presence of self-sperm in both wild-type (N2) and *spe-40(it133)* mutant hermaphrodites (Fig 2A and 2C). However, we continue to observe sperm in the spermatheca of *spe-40(it133)* hermaphrodites even at day four of adulthood when wild-type have fully depleted their self-sperm (Fig 2B and 2D). In addition, we observe endoreduplication in the eggs in the uteri of the *spe-40(it133)* hermaphrodites even in day 1 adults. This phenomenon is not observed in the wild-type adults until day 4 of adulthood when all of the sperm have been depleted. These observations indicate that sperm from *spe-40* mutants do not fertilize the egg, despite being present and making contact with the egg as the egg passes through the spermatheca and into the uterus.

**Figure 2:**
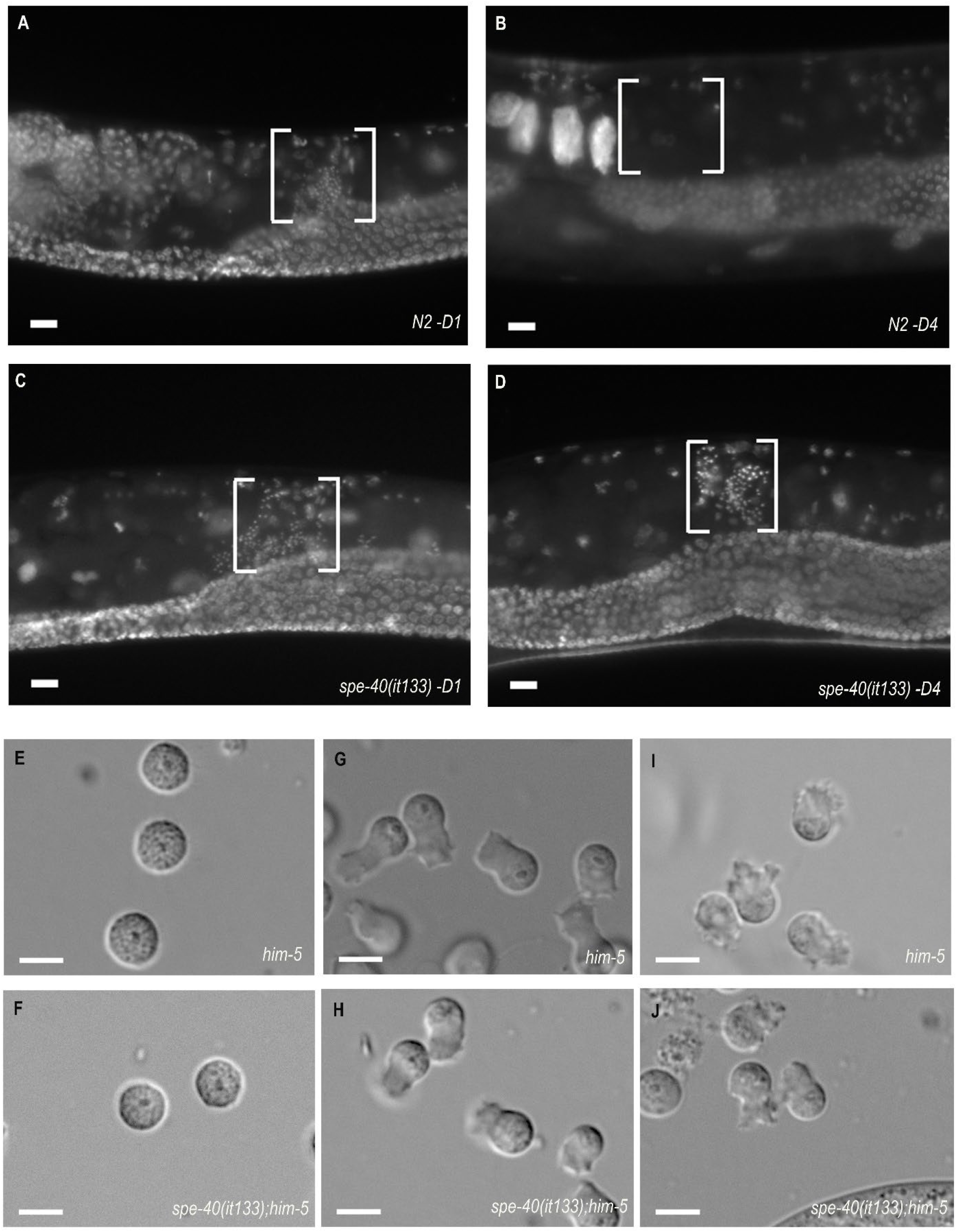
Sperm from *spe-40* mutants have normal morphology and activation but do not fertilize. **(A-D)** DAPI staining of wild-type and *spe-40(it133)* hermaphrodites at day one (D1) and day four (D4) of adulthood. The spermatheca is indicated by brackets and sperm nuclei are the small bright puncta within the spermathecae. The oviduct is to the right of the spermatheca and the uterus is to the left of the spermatheca. Scale bar = 12.5 µm **(E-F)** Unactivated spermatids dissected from control (*him-5*) and *spe-40(it133); him-5* males. Scale bar = 5 µm **(G-H)** *in vitro* (Pronase) activated spermatozoa from control (*him-5*) and *spe-40(it133); him-5* males. Scale bar = 5 µm **(I-J):** *In vivo* activated spermatozoa from control (*him-5*) and *spe-40(it133); him-5* males. Scale bar = 5 µm

Because sperm are present but fail to fertilize, we next examined sperm activation and morphology in *spe-40* mutants. We observed that spermatids from *spe-40(it133)* mutants are round with a prominent nucleus in the center indicating normal morphology that is indistinguishable from controls (Fig 2E-F). Next, we examined whether *spe-40(it133)* spermatids can undergo spermiogenesis/sperm activation. In *C. elegans*, spermiogenesis includes development of a pseudopod and acquisition of motility.^21^ Membranous Organelles (MOs) also fuse with the sperm plasma membrane and fertilization specific proteins relocate from the MOs to the sperm surface.^4,10,11,22–24^ *C. elegans* spermatids can be activated *in vitro* by exposure to Pronase.^21^ Sperm from *spe-40(it133)* mutant males are capable of activation *in vitro*, as indicated by formation of a pseudopod and normal morphology that is again indistinguishable from wild-type (Fig 2G-H). We also confirmed that the sperm from *spe-40(it133)* males activate *in vivo* by mating males with *fog-2* hermaphrodites and dissecting from the spermatheca the sperm that had been transferred to the hermaphrodite by the male (Fig 2I-J). We further analyzed the ultrastructure of sperm from *spe-40(it133)* mutants using Transmission Electron Microscopy (TEM). We found that they have normal morphology comprised of a single nucleus, pseudopod, fused MOs, and mitochondria (Fig 3). Despite defects in sperm function, sperm from *spe-40(it133)* mutants show no morphological defects. Therefore, we conclude that the reason sperm from *spe-40* mutants fail to fertilize the egg is because SPE-40 is specifically required for the fertilization process.

**Figure 3:**
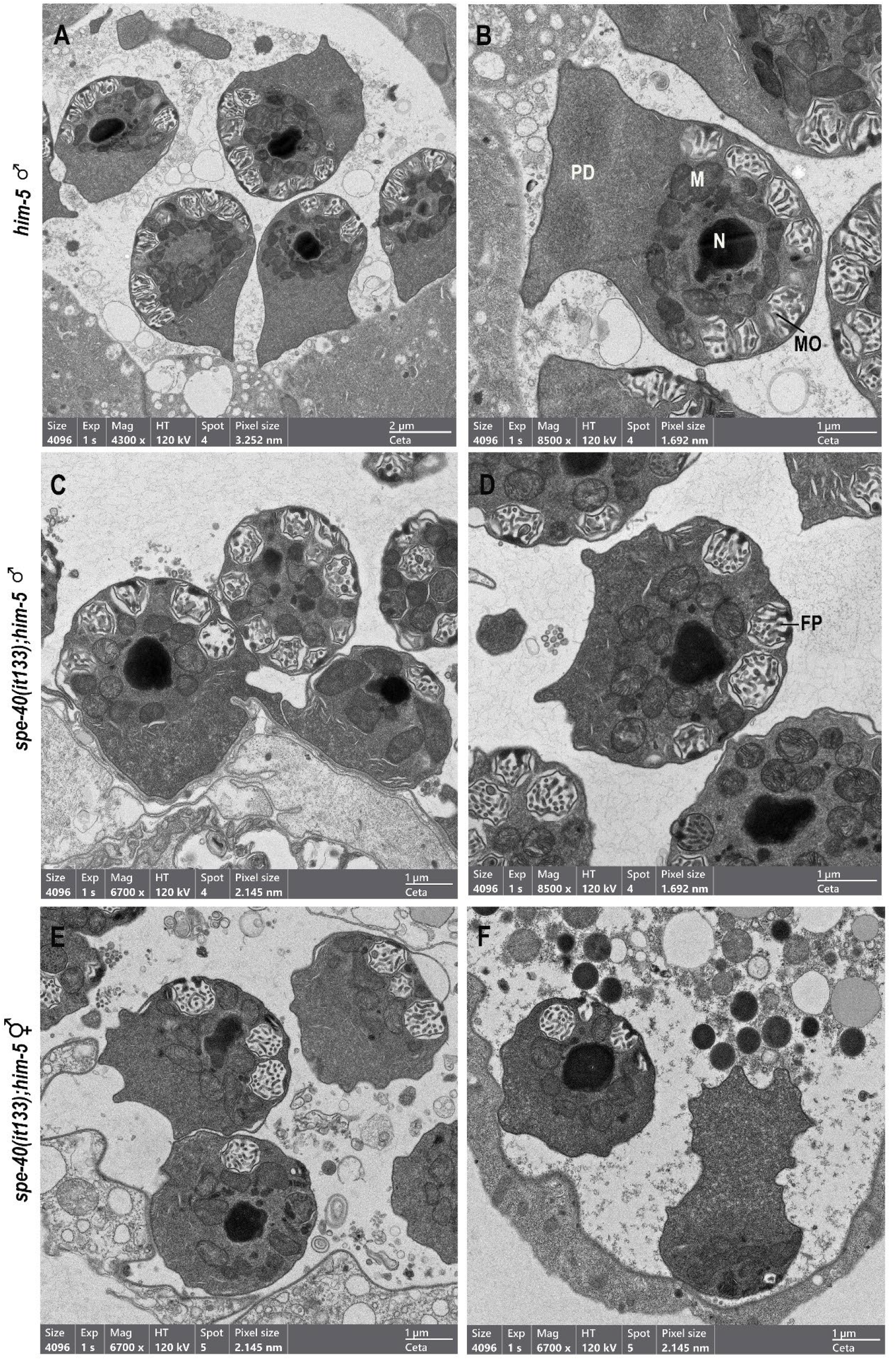
Transmission Electron Microscopy of spermatozoa shows sperm from *spe-40* mutants have no ultrastructural defects. **(A-D)** Males of the indicated genotype (*him-5* or *spe-40(it133); him-5)* were mated to *fog-2* hermaphrodites. The spermathecae of the hermaphrodites were then dissected to observe the sperm that had been transferred to the hermaphrodite by the male. Nucleus (N), Mitochondria (M), Pseudopod (P), Membranous Organelles (MOs) and Fusion Pore (FP). **(E-F)** The spermathecae of unmated *spe-40(it133); him-5* hermaphrodites were dissected to observe hermaphrodite self-sperm.

### SPE-40 is a transmembrane protein with an Ig domain

To identify the mutation responsible for the *spe-40* phenotype, we performed whole genome sequencing. Through a mapping by sequencing strategy^25^, we identified a mutation at the start codon of the ORF *Y37E11AR*.*7* positioned on Chromosome IV (3752583 – 3755455)^26^ in the *spe-40(it133)* strain. A single nucleotide change from C to T in the non-template sequence of *Y37E11AR*.*7* in the *spe-40(it133)* mutant causes a loss of the start codon and is predicted to result in no protein expression. We performed rescue experiments by creating extrachromosomal array lines carrying the *Y37E11AR*.*7* genomic sequence. Two independent *Y37E11AR*.*7* extrachromosomal arrays were able to rescue the fertility of *spe-40(it133)* back to wild-type levels (Fig 1A and S1A). To further confirm that we had identified the *spe-40* gene, we used CRISPR to create a second allele of *spe-40* where we inserted a premature stop codon and frameshift mutation at the end of Exon 1 of *Y37E11AR*.*7*. Brood sizing of *spe-40(nwk5)* hermaphrodites showed a severely sub fertile phenotype similar to the *spe-40(it133)* allele (Fig S1B). The SPE-40 protein is predicted to have a single transmembrane domain (DeepTMHMM^27^, SignalP^28^) and an Ig-like domain and is homologous to FAM187A/B in vertebrates (InterPro^29^).

### SPE-40 localizes to the cell body and pseudopod in spermatozoa

To observe the localization of SPE-40 in sperm, we used CRISPR to add an eGFP tag to the carboxy terminus of *spe-40* at the endogenous locus. In spermatids, SPE-40::eGFP is observed as bright puncta consistent with the location of MOs^5^ (Fig 4A-B). In *in vivo* activated sperm from *spe-40(nwk17[spe-40::egfp]); him-5* males, some SPE-40::eGFP is still in the MOs, but the protein has also relocated to the plasma membrane and is seen on the cell body as well as the pseudopod (Fig 4C-D). This matches the localization of other *C. elegans* sperm fertilization proteins^10,11,24^ and places SPE-40 in a position to directly interact with the surface of the egg.

**Figure 4:**
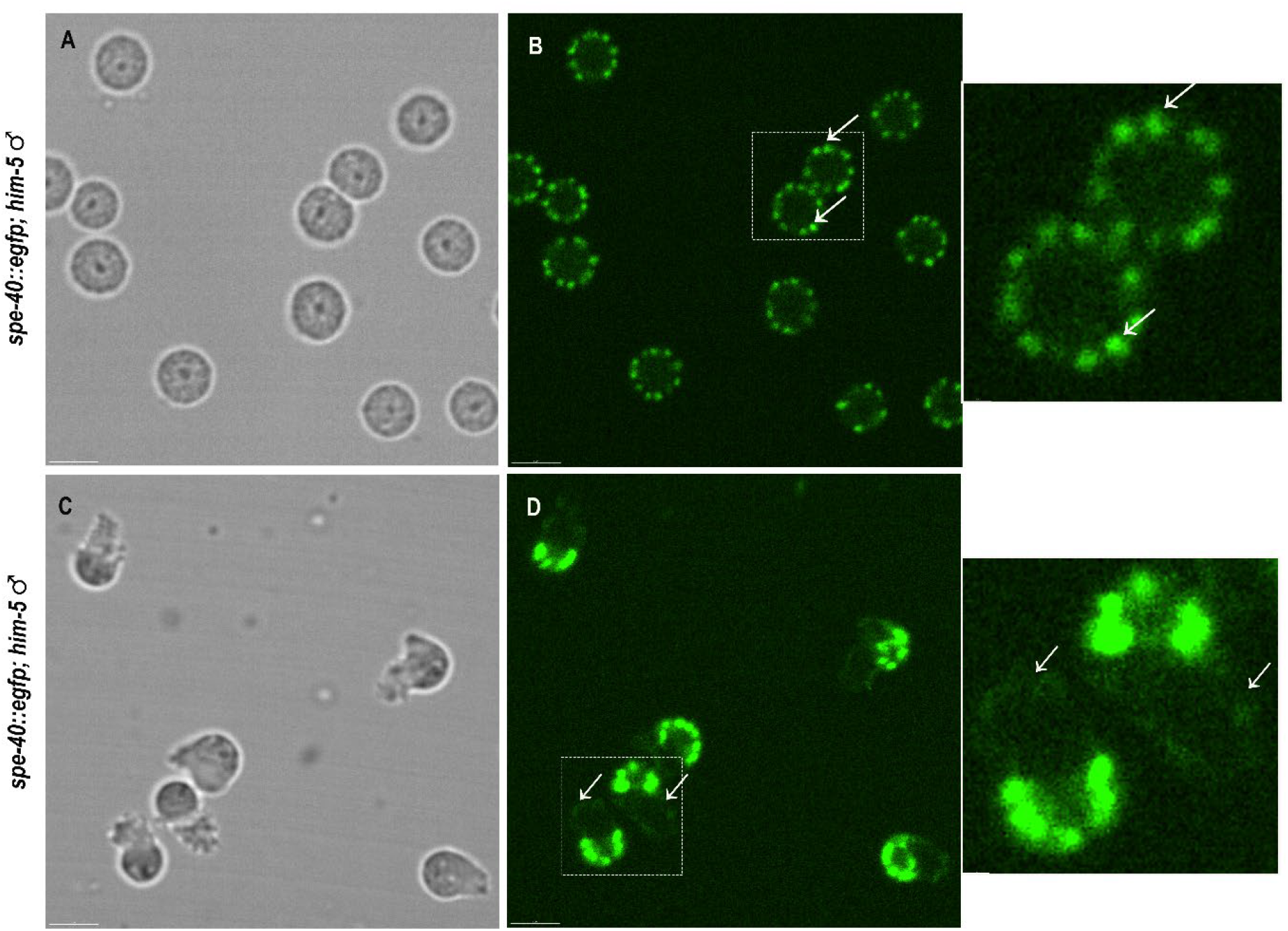
SPE-40 moves from the MOs to the cell surface after sperm activation. **(A-B)** SPE-40::eGFP in unactivated spermatids. The protein is concentrated in puncta reminiscent of MOs (white arrows). Scale bar = 5 µm **(C-D)** SPE-40::eGFP in *in vivo* activated spermatozoa. The protein is associated with the pseudopod (white arrows) as well as the cell body. Scale bar = 5 µm

## DISCUSSION

Through this work, we have shown that SPE-40 functions specifically during fertilization. Mutations in *spe-40* result in severe sub-fertility due to a sperm-specific defect. Importantly, we found that *spe-40* mutants have normal sperm morphology, sperm activation, competition, and retain their ability to migrate to and maintain their position at the site of fertilization. However, sperm from *spe-40* mutants do not fertilize.

SPE-40 is a single pass type I transmembrane protein with a single Ig-like domain. The SPE-40 protein is almost exclusively extracellular, consistent with a role in recognition, adhesion, and/or fusion with the egg membrane. In addition to SPE-40, there are nine other sperm proteins reported to be required for fertilization in *C. elegans*.^3–12^ Loss of any one of these proteins results in complete sterility or severe subfertility. When these proteins have been endogenously tagged and observed in live cells they have all been found to localize to MOs in spermatids and to translocate to the cell body and pseudopod in activated fertilization-competent spermatozoa. This common localization pattern places them in a position to act in cis with one another as well as in trans with the egg surface. Considering the importance of fertilization for the continuation of all sexually reproducing species, it seems counter-intuitive that instead of finding redundancy, we have discovered ten individual proteins that are each absolutely required specifically for fertilization. This discovery is not unique to *C. elegans* as vertebrates now have a similar number of sperm fertilization proteins reported.^30^ This led us to the hypothesis that these proteins work together in a multi-protein complex on the sperm surface and suggests an interdependency among the proteins for the formation or function of this complex^31^. This hypothesis is now strongly supported by structural predictions and proteomic analyses.^30,32–34^

SPE-40 has orthologues in vertebrates including human, mouse, and zebrafish. Like *C. elegans*, zebrafish have a single ortholog while mammals, including human and mouse, have two paralogues: FAM187A and FAM187B. Male fish lacking FAM187 and male mice lacking FAM187A are sterile, consistent with this protein being required for fertilization in these species as well.^30^ This suggests a deeply conserved role for SPE-40/FAM187 in mediating fertilization. How this conserved factor interacts and functions with taxa-specific proteins to mediate recognition, adhesion, and fusion between gametes remains an open question.

Ig-like domains are present in several fertilization proteins^7,8,11,33–36^, including SPE-40/FAM187. The presence of Ig-like domains near the cell surface as a part of a transmembrane protein or a secreted protein provides a stable structural support while also promotes the homotypic and heterotypic interaction with other domains of the proximal proteins.^37,38^ Thus, we hypothesize that Ig-like domain containing proteins may help facilitate the formation of premade ligand clusters on the sperm surface, facilitating the rapid interaction with an egg receptor initiating the process of fertilization. Interestingly, in species where HAP2 has been identified as the gamete fusogen^39–44^, there are no clear orthologs of SPE-40/FAM187, suggesting the possible evolution of two different mechanisms of fertilization. Complete identification of the proteins that mediate fertilization and a mechanistic understanding of their functions across species will be critical to understanding this fundamental biological process.

Understanding the genetic underpinnings of infertility is important for proper diagnosis and selection of the most appropriate and effective assisted reproductive technologies and treatments. While infertility can have a number of underlying causes, including hormonal imbalances and defects in gametogenesis, it can also be caused by mutations in the proteins that directly mediate fertilization. In these cases, measurable parameters such as sperm count and morphology are unaffected, making it critical to know the identity of the specific genes and proteins. This study, and the complementary study in vertebrates^30^ establish SPE-40/FAM187 as a sperm protein required for fertilization, adding it to a surprisingly large and still expanding collection of proteins.

## METHODS

### C. elegans

*C. elegans* were cultured on MYOB (Modified Youngren’s, Only Bacto-peptone) plates seeded with OP50 *E. coli*. The general maintenance of *C. elegans* was performed as described previously.^45^ All results shown here are performed with *C. elegans* maintained at 20 °C. The *spe-40(it133)* mutation was isolated from an EMS (ethylmethane sulfonate) mutagenesis screen by Dr. Diane Shakes while in the Kempheus Laboratory at Cornell University. A full list of strains used in this study is provided in Supplementary Table 1.

### Hermaphrodite fertility assay

To assess hermaphrodite fertility, L4 stage hermaphrodites were isolated on freshly seeded MYOB plates with OP50 and transferred every day to a new plate until they began laying unfertilized oocytes. The number of progeny on each plate was counted after 72 hours at 20 °C and the total number of progeny for each worm was determined.

### Male fertility assay

To assess male fertility, *fog-2* feminized L4 stage hermaphrodites were crossed with *him-5* or *spe-40(it133); him-5* males in a ratio of 1:3 at 20 °C. The males on the plates were removed after 24 hours and hermaphrodites were transferred to a new plate. The hermaphrodites were transferred every day to a new plate until they started producing unfertilized oocytes. The number of progeny on each plate was counted after 72 hours at 20 °C and the total number of progeny produced by each hermaphrodite was determined.

### Sperm competition assay

To assess sperm competition, *dpy-5* hermaphrodites at the L4 stage were crossed with *him-5* or *spe-40(it133); him-5* males in a ratio of 1:3. After allowing them to mate for 24 hours, the males were removed and the hermaphrodites were transferred to new plate and allowed to lay eggs for an additional 24 hours. The number of wild-type and Dpy progeny on each plate was counted after 72 hours at 20 °C.

### DAPI Staining

Hermaphrodites were fixed with ice-cold methanol for 30 – 60 seconds followed by 1X PBS wash. PBS was replaced with 10 µl VectaShield Antifade Mounting Media containing DAPI (Vector Laboratories). The worms were moved to a thin agar pad on a glass slide and a coverslip was added. Worms were viewed with a Zeiss Axio Observer inverted microscope using a 20X objective.

### Sperm assessments

All dissections were performed in 1X Sperm Media (50mM HEPES, 25mM KCl, 45mM NaCl,1 mM MgSO4, 5mM CaCl2, pH adjusted to 7.8 using NaOH) with 10 mM Dextrose.^46^ To observe spermatids and *in vitro* activated sperm, *him-5* or *spe-40(it133); him-5* males were isolated the day before the dissection. For dissections, the males are placed in a small drop of Sperm Media on a Histobond Adhesive slide (VWR) and a single clean-cut dissection of the gonad was performed. To assess *in vitro* activation, the Sperm Media was prepared with Pronase at a final concentration of 20 µg/ml. A coverslip was placed on the samples and sperm were imaged with a Zeiss Axio Observer inverted microscope using a 63X objective.

To observe sperm that were activated *in vivo, fog-2* hermaphrodites were allowed to mate with *him-5* or *spe-40(it133); him-5* or *spe-40(nwk17[spe-40::egfp]); him-5* males for 24 hours. The mated hermaphrodites were then dissected to release the sperm into the sperm media. To assess *in vivo* activation, sperm were imaged with a Zeiss Axio Observer inverted microscope using a 63X objective. To assess SPE-40::eGFP localization, sperm were imaged using an Andor Dragonfly spinning disk confocal microscope with a 63X objective.

### Transmission Electron Microscopy (TEM)

To assess male-derived sperm, *him-5* or *spe-40(it133); him-5* males were plated with *fog-*2 hermaphrodites for 24 hours. Gonads of mated *fog-2* hermaphrodites and unmated *spe-40(it133); him-5* hermaphrodites were dissected in sperm media. The samples were fixed using freshly prepared 2% glutaraldehyde and 2% formaldehyde in 0.1M sodium cacodylate buffer, pH 7.4 and incubated for 1 hour at room temperature. After, 4% low melting agar was added on to the fixed sample and the sample was kept at 4 °C to solidify before being cut into blocks using a razor blade. Samples were washed with 0.1 M sodium cacodylate buffer, pH 7.4 and post-fixed with 1% osmium tetroxide for 2 hours at room temperature. Samples were stained *en bloc* overnight in 1% uranyl acetate (aq) at room temperature followed by dehydration in an increasing gradient of acetone followed by 95% acetone overnight. After being dehydrated again in anhydrous 100% glass-distilled acetone, samples were infiltrated with increasing gradient series of Spurr’s resin. The samples were embedded in aluminum dishes and polymerized at 60°C for 2 days. Images were collected using a Thermo Scientific Talos L120C Transmission Electron Microscope.

### Whole Genome Sequencing

To determine the causative mutation in the *spe-40(it133)* strain, we took a SNP-mapping by sequencing approach.^47^ *spe-40(it133)* hermaphrodites were crossed with the polymorphic wild-type Hawaiian strain CB4856. F2 progeny with the sterile *spe-40* phenotype were pooled and genomic DNA was isolated for whole genome sequencing following previously established protocols.^48^ Library preparation and sequencing was performed by the University of Delaware DNA Sequencing & Genotyping Center. Samples were run on a NextSeq 2000. The data was analyzed using MiModD (https://mimodd.readthedocs.io/en/latest/nacreousmap.html) on a public Galaxy server.

### Preparation of transgenic rescue lines

Young adult N2 hermaphrodites were microinjected with a PCR product of genomic *Y37E11AR*.*7* including 766 bp of DNA upstream and 1578 bp downstream of the coding sequence along with myo-3p::gfp (100 μg/mL) as a transformation marker. Once transgenic lines were established the extrachromosomal array was crossed into *spe-40(it133)* to establish rescue of the sterile phenotype.

### CRISPR editing

To create the *spe-40(nwk5)* and *spe-40(nwk17[spe-40::egfp])* alleles we performed CRISPR editing following the general protocol described in Paix et al.^49^ The *dpy-10* co-conversion method was used to screen for successful injection plates.^50^ crRNA guide RNAs and DNA repair templates were ordered from IDT. For the *spe-40(nwk5)* allele the crRNA sequence was GUACACUUAUUUGGCAAUUG and the repair template was ATGTGGCTATTTGGTTCCGTCGTACAAATATCACTACTGTACACTTATTTGGCAAGCTAGCTTTGCGGTAAATATTTGGGGGATTTTTTTGCTATACACGGGAGTTTATGGAAAAGTTATCAACATACAAAATCAAAAAAAAACATTTCCCAGTTGACTTATC which inserts the sequence (5’-GCTAGCT-3’) containing an NheI restriction site and frameshift after bp 55 of the coding sequence. For the *spe-40(nwk17[spe-40::egfp])* allele we used the nested CRISPR method.^51^ For Step 1, the crRNA sequence was GUACAACUGACUGGCAACCA and the repair template was <u>TTCAGCGCCTTCATTTTCATCTTATCTCTCGTTGCCAGTCAATTGTACCGC</u>TCCAAGGGA GAGGAGCTCTTCACCGGAGTCGTCCCAATCCTCGTCGAGCTCGACGGAGTCAAGGAGT TCGTCACCGCTGCCGGAATCACCCACGGAATGGACGAGCTCTACAAG<u>TAAAATTTTGCA GTTGAATGAAGAAATTCCCCCTA</u> (homology arms to *spe-40* are underlined). Worms were genotyped by PCR to confirm the presence of the edit. Once homozygous animals were established worms were sequenced by Sanger sequencing to confirm the complete insert was present and in the correct position. Flanking primers to genotype *spe-40(nwk5)* were CTTCTGACAGCTCGACATTT and GCCAACAACTTATCGATACG. Flanking primers to genotype *spe-40(nwk17)* were GCCCGATAAGCCCATTAAGA and GTGAGAGAGTGCGGGATAAC.

### Statistical analysis

GraphPad Prism software was used to graph results and perform statistical analysis. T-tests were used to determine statistical significance for brood sizing assays.

## Acknowledgements

We would like to thank Diane Shakes for the original discovery of *spe-40(it133)* and Andrea Pauli and Steven Tang for sharing their findings and preprints with us. We would especially like to thank Shannon Modla in the UD Bio-Imaging Center for TEM sample preparation, imaging, and training. We also thank Timothy Chaya and Sylvain Le Marchand in the UD Bio-Imaging Center for assistance with SPE-40::GFP imaging. The Bio-Imaging Andor Dragonfly was acquired with a shared instrumentation grant (S10 OD030321) and microscopy access was supported by grants from the NIH-NIGMS (P20 GM103446, P20 GM139760) and the State of Delaware. This work was supported by the University of Delaware Research Foundation through a seed grant to ARK and by the Institutional Development Award (IDeA) from the National Institute of Health’s National Institute of General Medical Sciences under grant number P20GM103446 through a Graduate Core Facility Award to JNE for TEM imaging and training and through a Research Project Award to ARK.

---

**Supplementary Figure 1:**
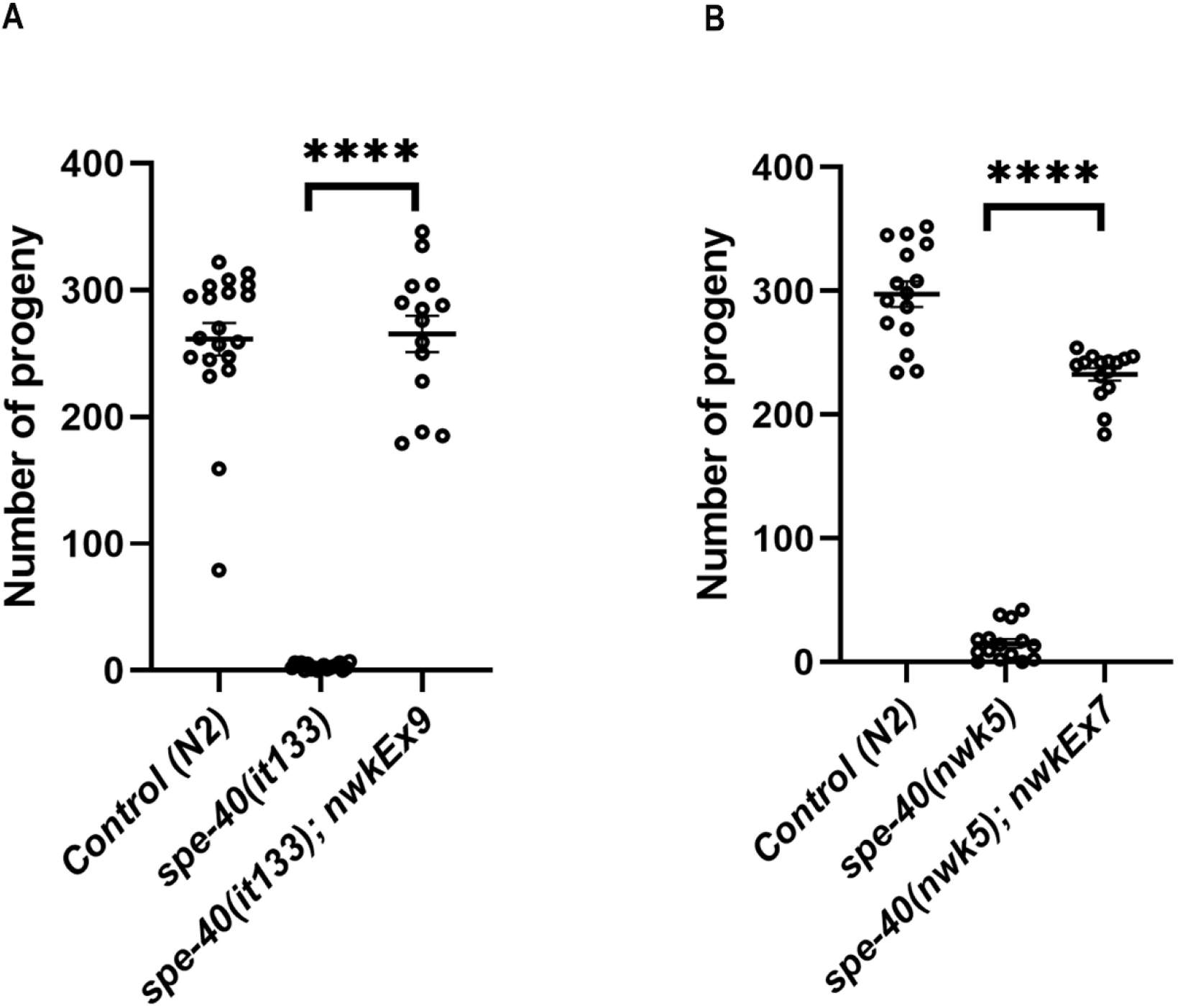
Brood sizes for additional strains, Related to Figure 1. **(A)** A second independent extrachromosomal array with the genomic *Y37E11AR*.*7* sequence rescues the brood size of *spe-40(it133)* hermaphrodites N≥15, p<0.0001 **(B)** Brood size of hermaphrodites for a second *spe-40* allele. *spe-40(nwk5)* was created by CRISPR and introduces a premature stop codon at the end of Exon 1. N≥15, p<0.0001

**Supplemental Table 1.**
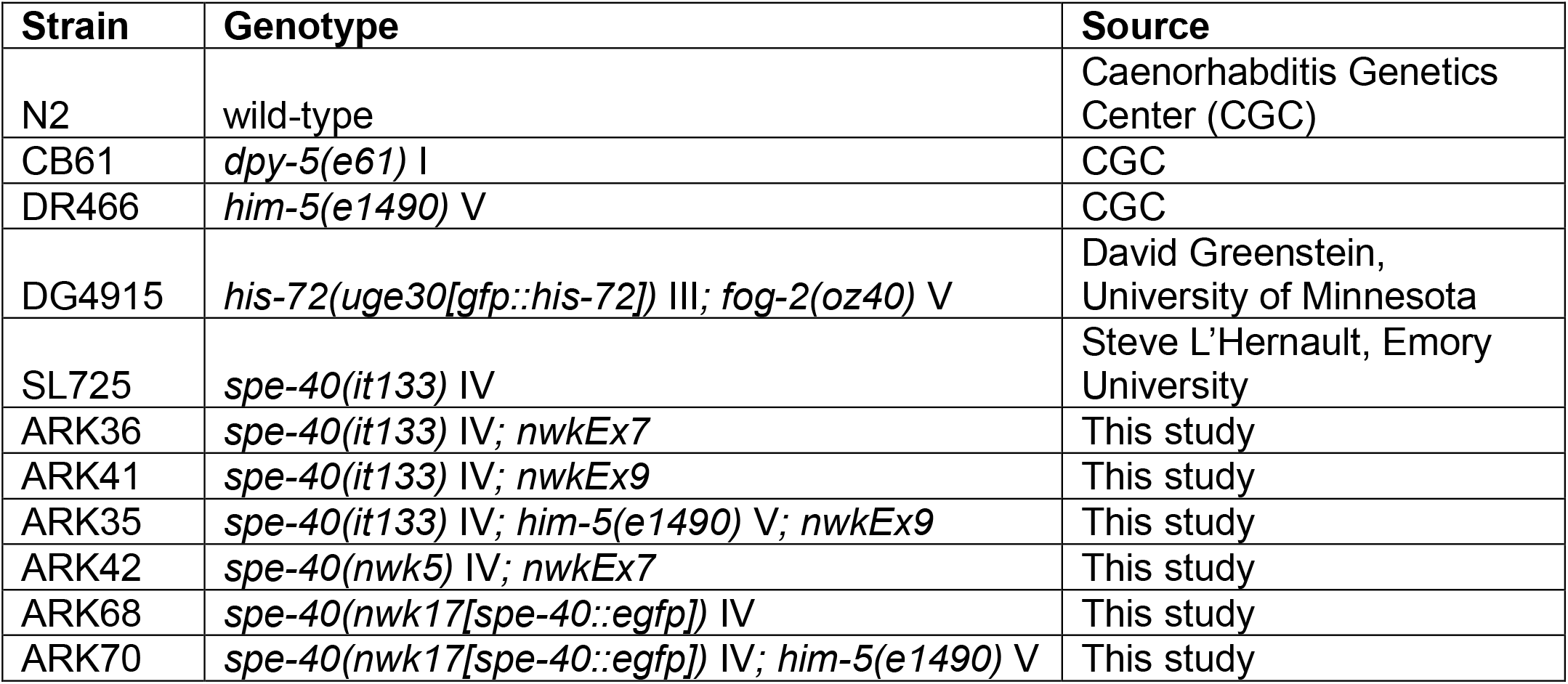
List of *C. elegans* strains used in this study.

